# MDS-causing mutations of ZRSR2 impair minor intron splicing by reducing its interaction with SF3B complex

**DOI:** 10.64898/2026.10.05.756190

**Authors:** Tomoki Chiba, Eri Matsumoto, Eiji Obayashi, So Masaki, Satoshi Tanaka, Naoyuki Kataoka

**Affiliations:** Laboratory of Cellular Biochemistry, Department of Animal Resource Sciences, Graduate School of Agricultural and Life Sciences, The University of Tokyo, Tokyo, Japan; Department of Agricultural Chemistry, Faculty of Applied Biosciences, Tokyo University of Agriculture, Tokyo, Japan; Department of Biochemistry, Shimane University School of Medicine, Izumo, Japan

## Abstract

Minor introns are rare but evolutionarily conserved introns in the human genome. Defects in minor spliceosome components have been reported to cause several human diseases. Mutations in ZRSR2 are frequently observed in myelodysplastic syndrome (MDS) patients. However, the precise molecular mechanism by which these mutations cause a loss of ZRSR2 function in minor intron splicing remain elusive. In this study, we characterized the impact of MDS-causing missense mutations in ZRSR2. Most of the mutant ZRSR2 proteins we tested showed different subnuclear localization pattern from that of wild-type ZRSR2 protein. Co-immunoprecipitation (co-IP) analysis revealed that UHM missense mutations disrupted the interaction between ZRSR2 and SF3B complex. We also demonstrated that ZRSR2 preferentially immunoprecipitated minor spliceosome components over major spliceosome components and it directly interacts with SF3B1. Furthermore, co-IP analysis utilizing a variety of ZRSR2 mutants revealed that mutations in the UHM and Zn2 domains significantly attenuated the interaction with SF3B complex. Finally, UHM and Zn2 mutations markedly disrupted the supporting activity of ZRSR2 for minor intron splicing. Our results indicate that disruption of the physical interaction with SF3B complex is one of the hallmarks of MDS-causing mutations of ZRSR2, which results in aberrant splicing of minor introns.

## Introduction

Pre-mRNA splicing is a fundamental process for gene expression in higher eukaryotes (Wahl et al. 2009). During splicing reaction, introns are excised and exons are ligated with high accuracy. Mutations of trans-acting factors or cis-regulatory RNA elements often result in various human diseases (Scotti and Swanson 2016; Anna and Monika 2018; Love et al. 2023; Norppa et al. 2025). One of such RNA splicing-deficient diseases is preleukemic cancer condition, myelodysplastic syndrome (MDS). In MDS, genomic mutations in hematopoietic stem cells (HSCs) disrupt their proliferation and differentiation, which leads to ineffective hematopoiesis (Dotson and Lebowicz 2026). Genomic-sequence analyses with MDS patient samples have revealed that mutations in splicing factors (SF3B1, U2AF1, SRSF2, and ZRSR2) are frequently detected (Yoshida et al. 2011; Haferlach et al. 2014), suggesting that incorrect splicing by splicing factor mutations is a cause of MDS. While MDS-causing mutations of SF3B1, U2AF1 and SRSF2 are concentrated at specific amino acid residues, diverse somatic mutations including missense, nonsense and frameshift mutations in ZRSR2 gene are reported across the entire coding region of ZRSR2 (Yoshida et al. 2011; Haferlach et al. 2014; Verma et al. 2018; Inoue et al. 2021; Norppa et al. 2025). These findings strongly suggest that ZRSR2 mutations in MDS cause ‘loss of function’ of ZRSR2.

There are two types of introns, U2-type and U12-type introns, in eukaryotic genome. U12-type introns are also referred as minor introns because they account for only 0.4% of all introns (Wu and Krainer 1999; Kataoka et al. 2021). Minor introns share consensus sequences distinct from those of major introns, and they are characterized by conserved 5’ splice site (5’ss), branch point sequence (BPS) and weak polypyrimidine tract (PPT) (Turunen et al. 2013; Verma et al. 2018; Kataoka et al. 2021; Norppa et al. 2025). These two types of introns are excised by two distinct splicing machineries. Pre-mRNA splicing is carried out by a large RNA-protein complex, termed spliceosome, which is composed of uridine-rich small nuclear RNAs (U snRNA) and diverse splicing factors (Wahl et al. 2009; Turunen et al. 2013; Verma et al. 2018; Kataoka et al. 2021; Norppa et al. 2025). The major spliceosome composed of U1, U2, U4/U6 and U5 small nuclear ribonucleoprotein (snRNP) function in excising major introns. In contrast, U11, U12, U4atac, U6atac and U5 snRNP are assembled to form minor spliceosome in the excision of minor introns. These two spliceosomes also possess common factors such as SF3B complex composed of SF3B1-B6. Splicing reaction begins with the recognition of the RNA elements in pre-mRNA (Wahl et al. 2009; Turunen et al. 2013; Kataoka et al. 2021). In major intron splicing, U1 and U2 snRNPs recognize 5’ss and BPS, respectively, and U2AF1/2 heterodimer also contacts strong PPT and 3’ splice site (3’ss). In contrast, U11/U12 di-snRNP recognizes 5’ss and BPS in minor intron splicing (Wahl et al. 2009; Turunen et al. 2013; Kataoka et al. 2021).

ZRSR2 is also referred to as U2AF1-related protein (URP) due to the amino-acids sequence homology with U2AF1, which has a function in the recognition for 3’ss of major intron (Tronchere et al. 1997; Merendino et al. 1999; Wu et al. 1999; Zorio and Blumenthal 1999; Shen et al. 2010). Like U2AF1, ZRSR2 harbors a U2AF homology motif (UHM), which is thought to interact with U2AF ligand motif (ULM), two Zinc Finger domains (Zn1 and Zn2) and Arginine and Serine-rich domain (RS) (Figure 1A)(Tronchere et al. 1997; Kielkopf et al. 2004; Loerch and Kielkopf 2016). In addition, ZRSR2 has N-terminal domain (N) and basic amino acids rich domain (BARD) that U2AF1 does not possess (Figure 1A). Missense mutations are observed in all domains of ZRSR2, suggesting that each domain has specific functions.

**Figure 1.**
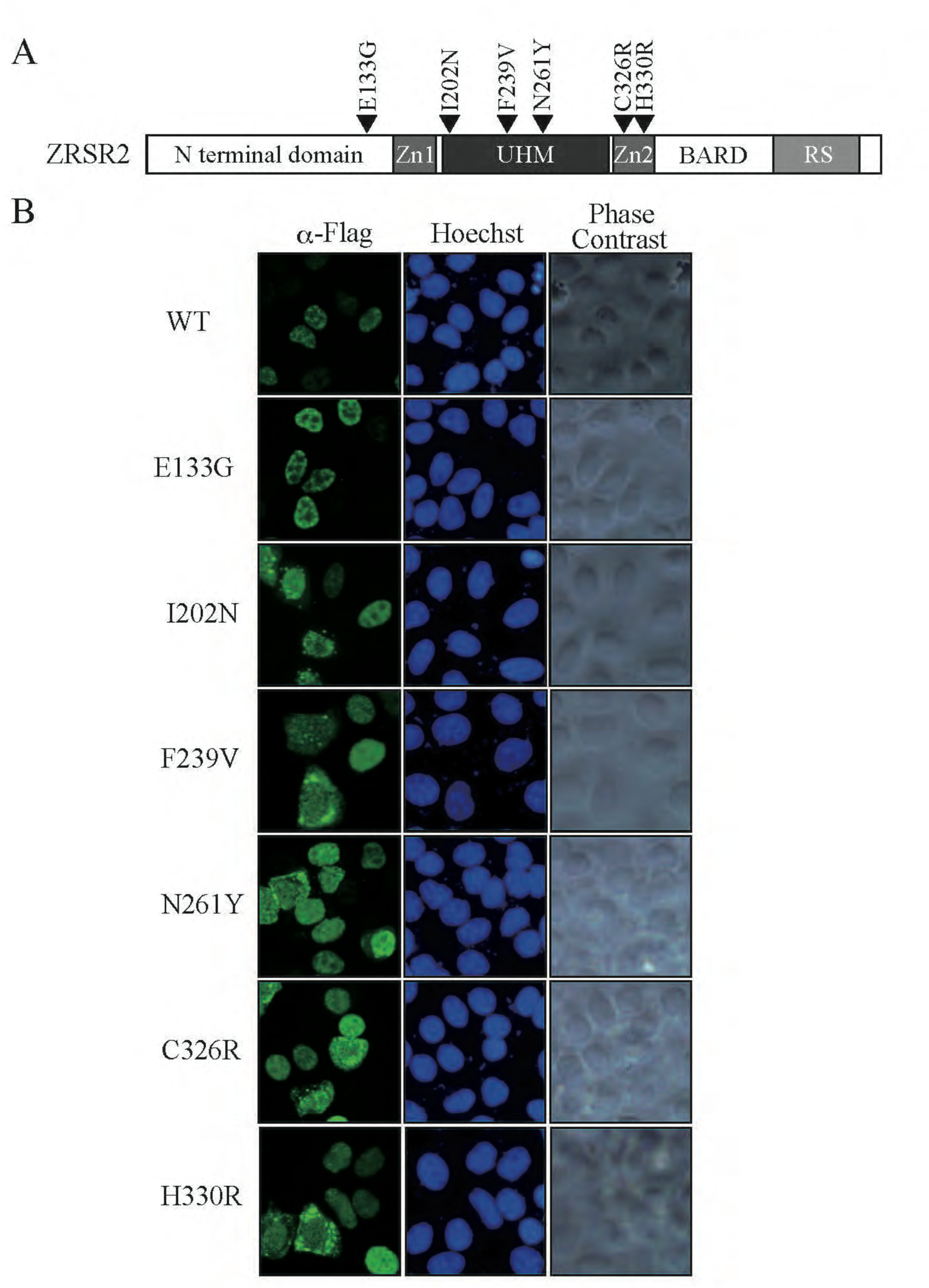
Subcellular localization of Flag-tagged wild-type (WT) and mutant ZRSR2 proteins in HeLa cells. (A) Schematic representation of ZRSR2 protein with domain structure. N terminal domain: amino-terminal domain, Zn1: Zinc Finger domain 1, UHM: U2AF homology motif, Zn2: Zinc Finger domain 2, BARD: Basic amino-acid rich domain, RS: Arginine-Serine rich domain. (B) Transfected HeLa cells were stained with anti-Flag antibody. WT panel shows subcellular localization of wild-type ZRSR2 protein. Other panels indicate the subcellular localization patterns of the indicated ZRSR2 mutant proteins. The panels Hoechst indicate the position of HeLa cell nuclei, and the Phase Contrast panels show the phase-contrast images of the cells. The scale bar indicates 20 µm.

By *in vitro* splicing assay, it was demonstrated that ZRSR2 was crosslinked with the 3’ss of both major and minor introns (Shen et al. 2010). In addition, it was suggested that ZRSR2 plays a role in the initial assembly of minor spliceosome (Shen et al. 2010). RNA-sequencing analyses of ZRSR2-mutated MDS patient samples and ZRSR2-deficient cells revealed that loss of ZRSR2 functions mainly resulted in aberrant splicing (especially intron retention) of minor introns, while it appeared to have less impact on major intron splicing (Madan et al. 2015; Inoue et al. 2021). Also, the cross-linking and immunoprecipitation (CLIP) analysis indicated that ZRSR2 preferentially contacted U12 snRNA and minor introns (Kwon et al. 2024). Furthermore, a recent reconstruction assay of the minor spliceosome followed by Liquid Chromatography-Mass Spectrometry (LC-MS) analysis revealed that ZRSR2 was detected in the early-stage complex (pre-B complex) of minor intron splicing (Bai et al. 2024). Taken together it is assumed that ZRSR2 mainly has functions in minor splicing to assemble minor spliceosome via 3’ss recognition. However, molecular functions of multiple domains of ZRSR2 in minor intron splicing are still unclear.

In this study, we investigated how ZRSR2 mutations found in MDS patients affect its function. By utilizing both wild-type and mutant ZRSR2 proteins, we identified SF3B complex components as specific interacting partners of ZRSR2. We also demonstrated that ZRSR2 preferentially associate with U12 snRNP likely through physical interaction with U11/12 65K protein. Through comprehensive analysis of missense mutations, we show that missense mutations in UHM and Zn2 domains significantly disrupt the ZRSR2-SF3B interaction. Finally, we demonstrate that those UHM and Zn2 mutations markedly attenuate minor intron splicing ability of ZRSR2. Together, these results indicate that disruption of the ZRSR2-SF3B interaction, which highly likely lead to aberrant splicing of minor introns, is one of the hallmarks of MDS-causing mutations in ZRSR2.

## Results

### ZRSR2 proteins harboring MDS-causing mutations show aberrant localization patterns in HeLa cell nuclei

Many missense mutations in coding sequences were identified in the *ZRSR2* gene in MDS patients. To investigate the impact of MDS-causing mutations on the functions of ZRSR2, we first determined subcellular localization patterns of ZRSR2 mutants. We selected the representative mutations initially identified by Yoshida et al (Yoshida et al. 2011) (Figure 1A). We transfected HeLa cells with the plasmids expressing either wild-type or mutant C-terminal Flag-tagged ZRSR2 proteins and performed immunofluorescent staining utilizing anti-Flag tag antibody. As shown in Figure 1B, wild-type ZRSR2 protein showed speckle-like localization patterns in HeLa cell nuclei. In contrast, all the mutants except E133G exhibited different localization patterns from that of wild-type ZRSR2 in the nuclei.

### Identification of SF3B complex proteins as specific interactors of ZRSR2

The results shown in Figure 1 raised the possibility that MDS-causing mutations affected the interaction between ZRSR2 and its specific binding factors, which resulted in different subnuclear localization. To test whether such specific binding proteins exist or not, we carried out co-immunoprecipitation (co-IP) experiments using transiently expressed Flag-tagged ZRSR2 wild-type and mutant proteins (I202N, F239V). Cell lysates from Flag vector-transfected cells were used as a negative control. As shown in Figure 2A, several proteins showed different amount between fractions of the wild-type and two mutant proteins. Mass spectrometry analyses revealed that two proteins enriched in the wild-type co-precipitated fraction were SF3B1 and SF3B3, while two prominent proteins in the precipitated fraction with mutant proteins were hsp70 and hsp90 (Figure 2A). We then validated co-immunoprecipitation of SF3B complex proteins with ZRSR2 by co-IP experiments. Results shown in Figure 2B indicate that SF3B1 and SF3B3 proteins are efficiently co-precipitated with wild-type ZRSR2. In contrast, co-precipitation efficiencies of SF3B1 and SF3B3 proteins were significantly reduced in both I202N and F239V mutants (Figure 2B, lanes 7 and 8). We also determined whether another SF3B component, SF3B4 was co-precipitated or not. As shown in Figure 2B, SF3B4 was also efficiently co-precipitated with wild-type ZRSR2 but inefficiently with I202N and F239V mutant proteins. These results indicate that ZRSR2 associates with SF3B complex and that the MDS-causing I202N and F239V mutations markedly reduce this interaction.

**Figure 2.**
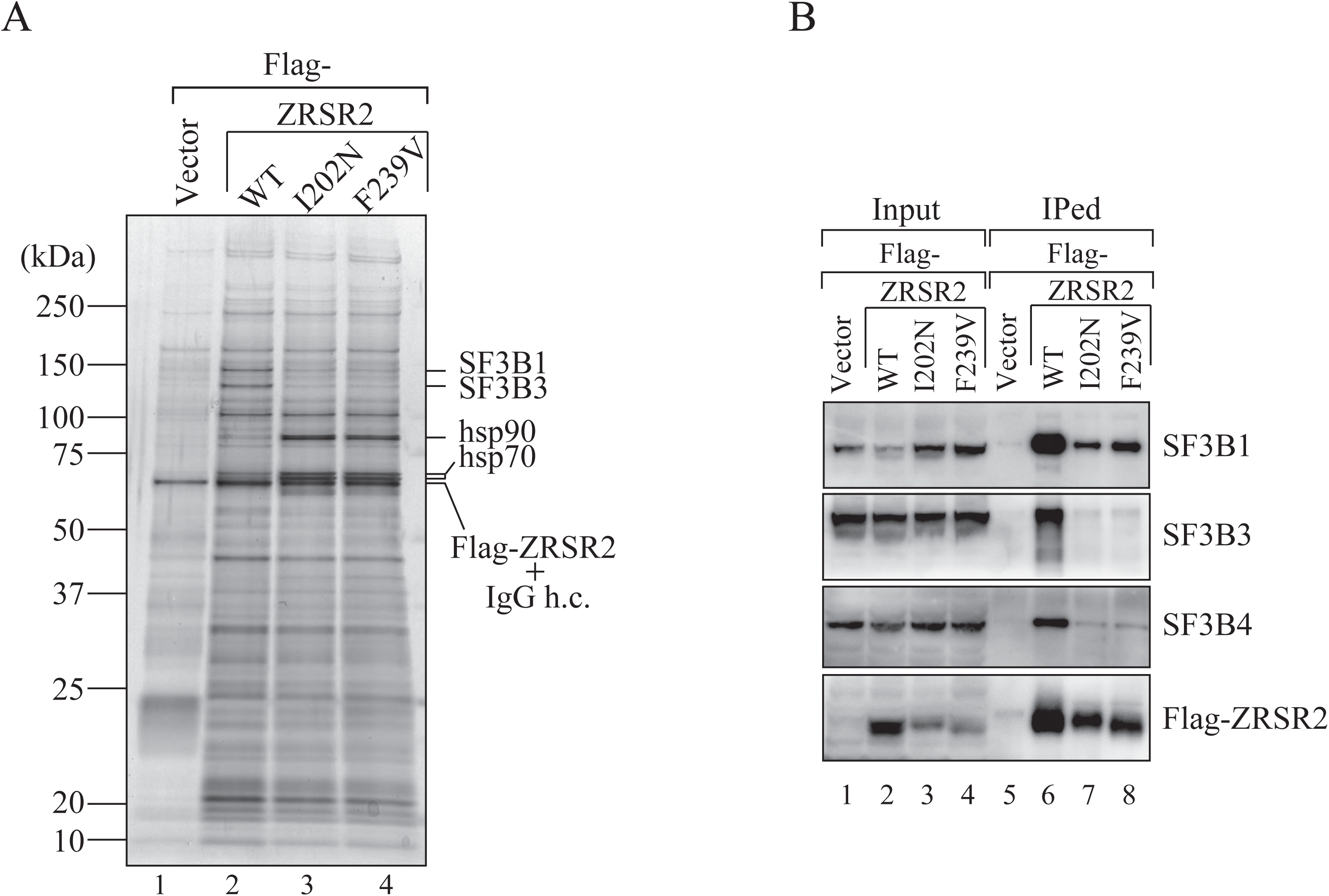
Analysis of co-immunoprecipitated proteins with wild-type and mutant ZRSR2 proteins. (A) Either Flag-tagged wild-type or two mutant ZRSR2 proteins were expressed in HEK293T cells and immunoprecipitated by anti-Flag M2 agarose. Co-precipitated proteins were analyzed by SDS-PAGE, followed by mass spec analysis to identify specific binders. The positions of specific binders are indicated with their names at the right of the panel. The positions of Flag-ZRSR2 and immunoglobulin G heavy chain (IgG h.c.) are also shown. (B) Validation of co-immunoprecipitation of SF3B proteins with ZRSR2 protein. Either Flag-tagged wild-type or two mutant ZRSR2 proteins were expressed in HEK293T cells and immunoprecipitated by anti-Flag M2 agarose. Precipitated proteins were analyzed by immunoblotting with specific antibodies indicated.

### ZRSR2 interacts with U12 spliceosome protein and SF3B complex proteins in an RNA-independent manner

SF3B complex is known to be recruited to both major and minor spliceosome and plays a crucial role in branch point recognition (Will et al. 2001). On the other hand, another SF3 component, SF3A complex, incorporates into major spliceosome but not into minor spliceosome (Will et al. 2004; Turunen et al. 2013; Zhao et al. 2025). To examine which type of spliceosome proteins that ZRSR2 preferentially interacts with, we performed co-IP analyses followed by immunoblotting with the antibodies against components of SF3A and SF3B complex and unique factors involved in each spliceosome. As shown in Figure 3A, SF3B components, SF3B1, SF3B3 were efficiently co-precipitated with wild-type ZRSR2. In addition, U11/12 snRNP 65K, a minor spliceosome component (Will et al. 2004; Benecke et al. 2005; Turunen et al. 2013; Zhao et al. 2025), also co-precipitated efficiently with ZRSR2 (Figure 3A, lane 4). However, co-precipitation of SF3A1 and U2 snRNP A, which are involved in major spliceosome (Turunen et al. 2013), was barely detected. These results demonstrate that ZRSR2 preferentially interacts with minor spliceosome components. We then further examined whether these interactions were RNA-mediated or not by the immunoprecipitation experiments both in the presence and absence of RNase A. U12 snRNA was effectively precipitated with ZRSR2 protein in the absence of RNase A (Figure 3B, lane 5), indicating that ZRSR2 protein associates with U12 snRNP. On the other hand, co-precipitation of U12 snRNA disappeared in the presence of RNase A, which indicates that RNAs including U12 snRNAs in cell lysates were degraded by the RNase addition (Figure 3B, lane 6). Co-precipitation of SF3B1, SF3B3 and U11/12 65K was observed even in the presence of RNase A, as well as in the absence of RNase (Figure 3B, lanes 5 and 6). These results indicate that ZRSR2 associates with SF3B proteins and U11/12 65K in an RNA-independent manner, most likely through protein-protein interaction.

**Figure 3.**
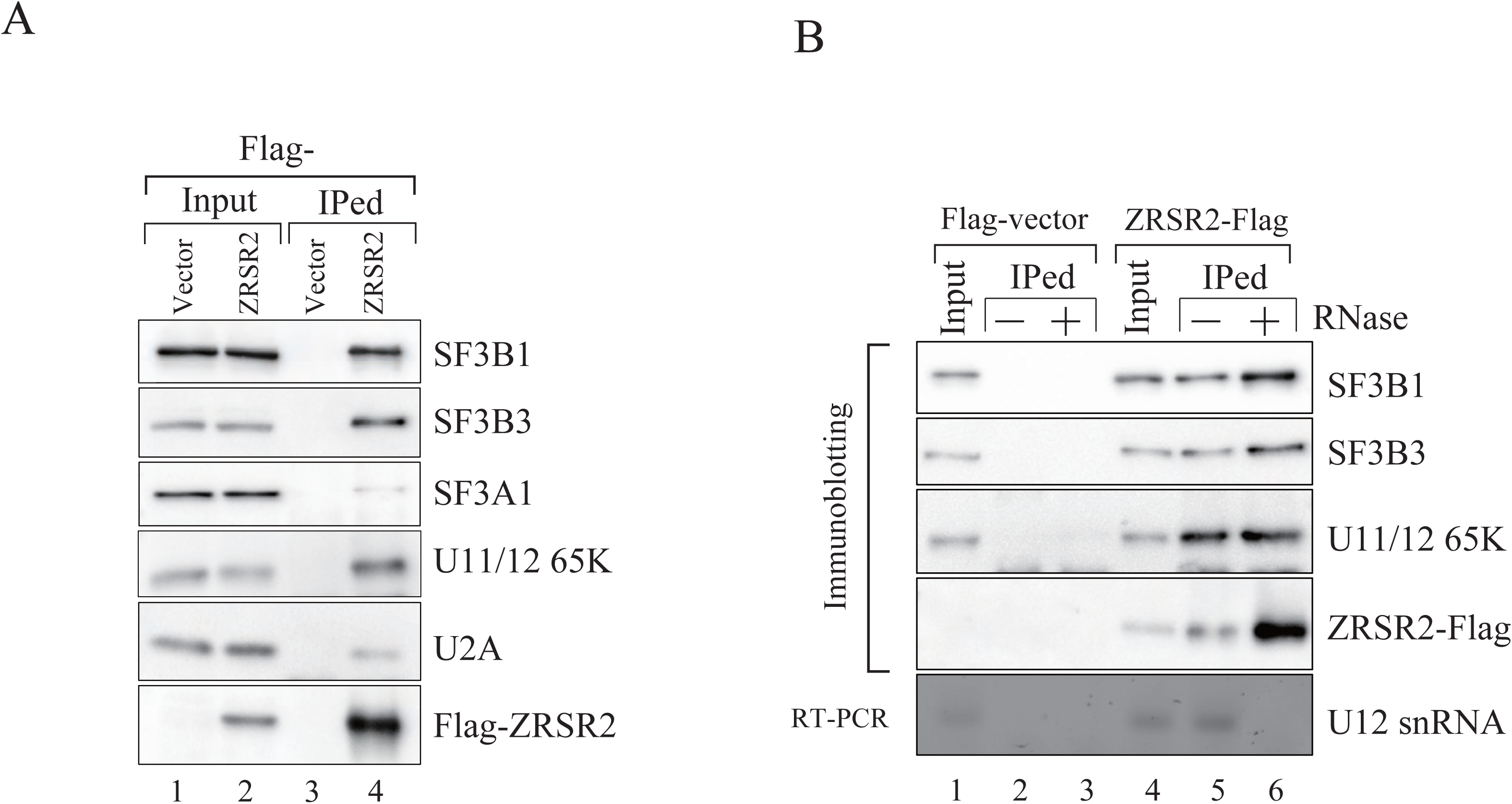
ZRSR2 protein interacts with SF3B complex and U12 snRNP in an RNA independent manner. (A) Flag-tagged ZRSR2 proteins were expressed in HEK293T cells and immunoprecipitated by anti-Flag M2 agarose. Precipitated proteins were analyzed by immunoblotting with specific antibodies as indicated. (B) Co-immunoprecipitation analysis in the presence or absence of RNase. Flag-tagged ZRSR2 proteins were expressed in HEK293T cells and immunoprecipitated by anti-Flag M2 agarose in the presence or absence of RNase A. Precipitated proteins were analyzed by immunoblotting with specific antibodies as indicated.

Since ZRSR2 possesses UHM, an attempt was made to determine whether this UHM interacts directly with U2AF ligand motif (ULM) of SF3B1 or not. To test this hypothesis, we prepared His-tagged ZRSR2 and GST-fused SF3B1 recombinant proteins by expressing them in *E. coli* cells and performed GST pull-down assay utilizing these recombinant proteins. As a bait, we first used the GST-tagged the amino-acid 330-394 region of SF3B1, which contains ULM5. As for preys, His-tagged ZRSR2 fragments spanning 1-356 or 151-356 was used as a prey. His-ZRSR2 151-356 contains Zn1, UHM and Zn2 domains of ZRSR2 (Figure 4A). GST-SF3B1 330-394 was able to pull down both His-tagged ZRSR2 proteins (Figure 4B, Lanes 4-6). Next, we carried out same pull-down experiments using either the former half (330-349) or the latter half (348-394) fragment of GST-SF3B1 330-394 protein. We found that the former half pulled down His-tagged ZRSR2 proteins, whereas the latter half did not (Figure 4, lanes 7-12). These results indicate that ULM5 in SF3B1 directly interacts with ZRSR2 through UHM-ULM interaction.

**Figure 4.**
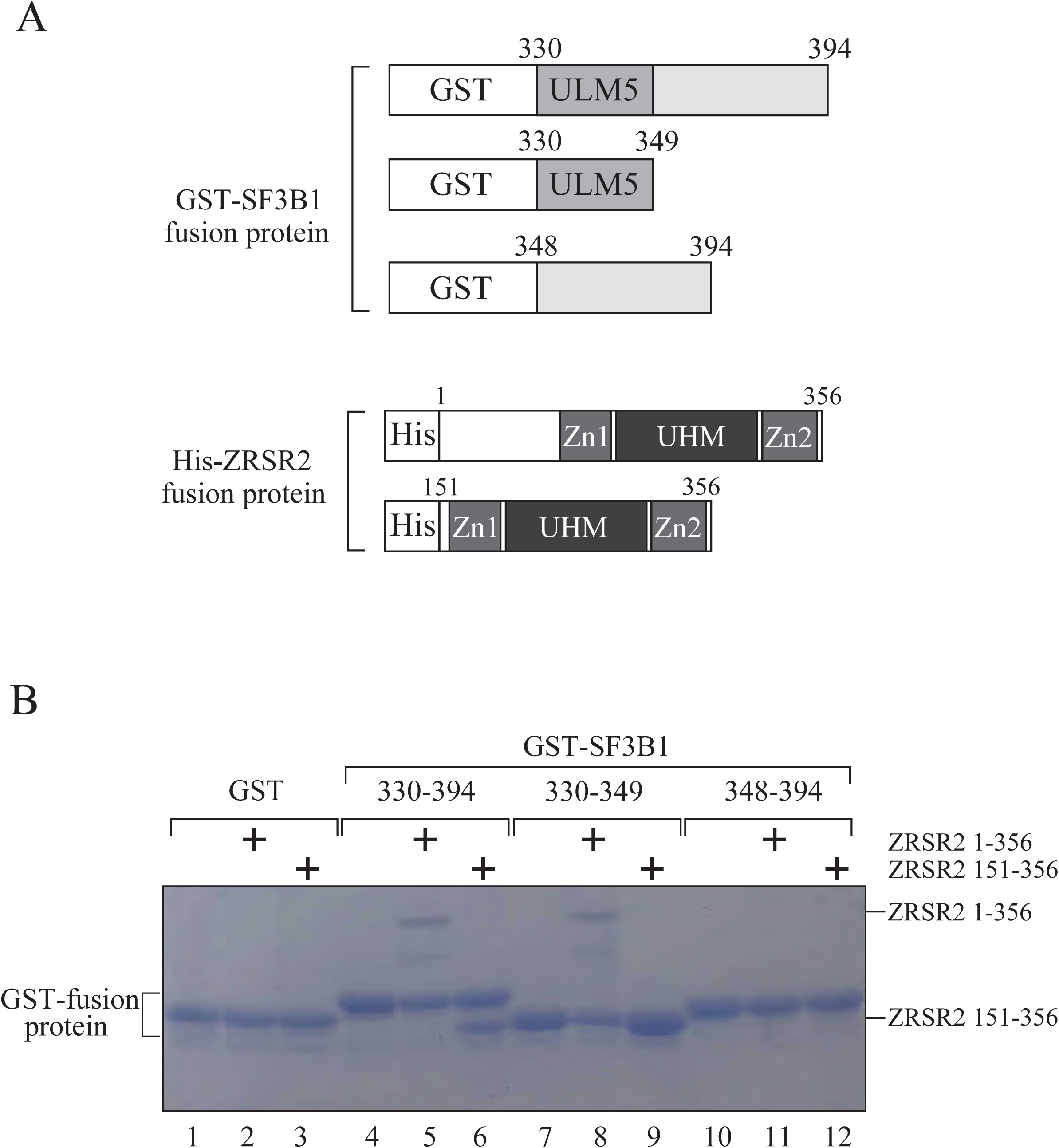
GST-pull down assay utilizing recombinant His-ZRSR2 and GST-SF3B1 protein. (A) Schematic representation of the recombinant proteins used for binding assays. GST : Glutathione S-transferase-tag, ULM5 : U2AF ligand motif 5, His : His-tag, Zn1 : Zinc Finger domain 1, UHM : U2AF homology motif, Zn2 : Zinc Finger domain 2 (B) GST-SF3B1 and His-ZRSR2 proteins were expressed in E. coli cells and purified. Those proteins were mixed with the indicated combination, and their binding was analyzed by SDS-PAGE. The positions of ZRSR2 proteins are shown at the right of the panel.

### UHM and Zn1/2 mutations in ZRSR2 drastically reduce its interaction activity with SF3B proteins

To date many of the mutations in ZRSR2 gene coding region have been identified in MDS patients (Verma et al. 2018; Norppa et al. 2025). Given our findings that mutations in the UHM of ZRSR2 impair SF3B-ZRSR2 interaction, we performed similar co-IP experiments to determine whether other ZRSR2 mutations affect the SF3B-ZRSR2 interaction or not. We prepared a series of ZRSR2 mutants that harbor a point mutation in each domain. First, we tested mutant ZRSR2 proteins harboring mutations in either N-terminus or Zn1 domain. The results in Figure 5A demonstrated that mutations in the N-terminal domain (G41R, E133D and E133G) had almost no effect on co-immunoprecipitation of SF3B proteins (lanes 10-12). In contrast, mutations in the Zn1 domain (C181S and H191Y) reduced SF3B protein co-immunoprecipitation (Figure 5A, lanes 13 and 14). Interestingly, none of the mutations affected co-immunoprecipitation of U12 snRNA and U11/U12 65K (Figure 5A, lanes 9-14). Next, we utilized mutant ZRSR2 proteins harboring mutations in UHM and Zn2. Like the results with I202N and F239V mutants, another UHM mutation, N261Y, also greatly reduced the interaction with SF3B proteins (Figure 5A, lane 12). Two Zn2 mutations (C326R and H330R) also exhibited impaired co-immunoprecipitation of SF3B proteins (Figure 5B, lanes 13 and 14). Neither the UHM nor the Zn2 mutants affected U12 snRNA and U11/U12 65K protein co-immunoprecipitation (Figure 5B, lanes 10-14). Lastly, we tested ZRSR2 mutants harboring mutations in the BARD or RS domain. None of the mutants exhibited reduction of co-immunoprecipitation efficiency for SF3B proteins (Figure 5C, lanes 10-14). These mutants also precipitated U12 snRNA and U11/U12 65K protein as efficiently as wild-type ZRSR2 (Figure 5C, lanes 10-14). Taken together, point mutations in Zn1, UHM and Zn2 domains of ZRSR2 impair interaction with SF3B complex.

**Figure 5.**
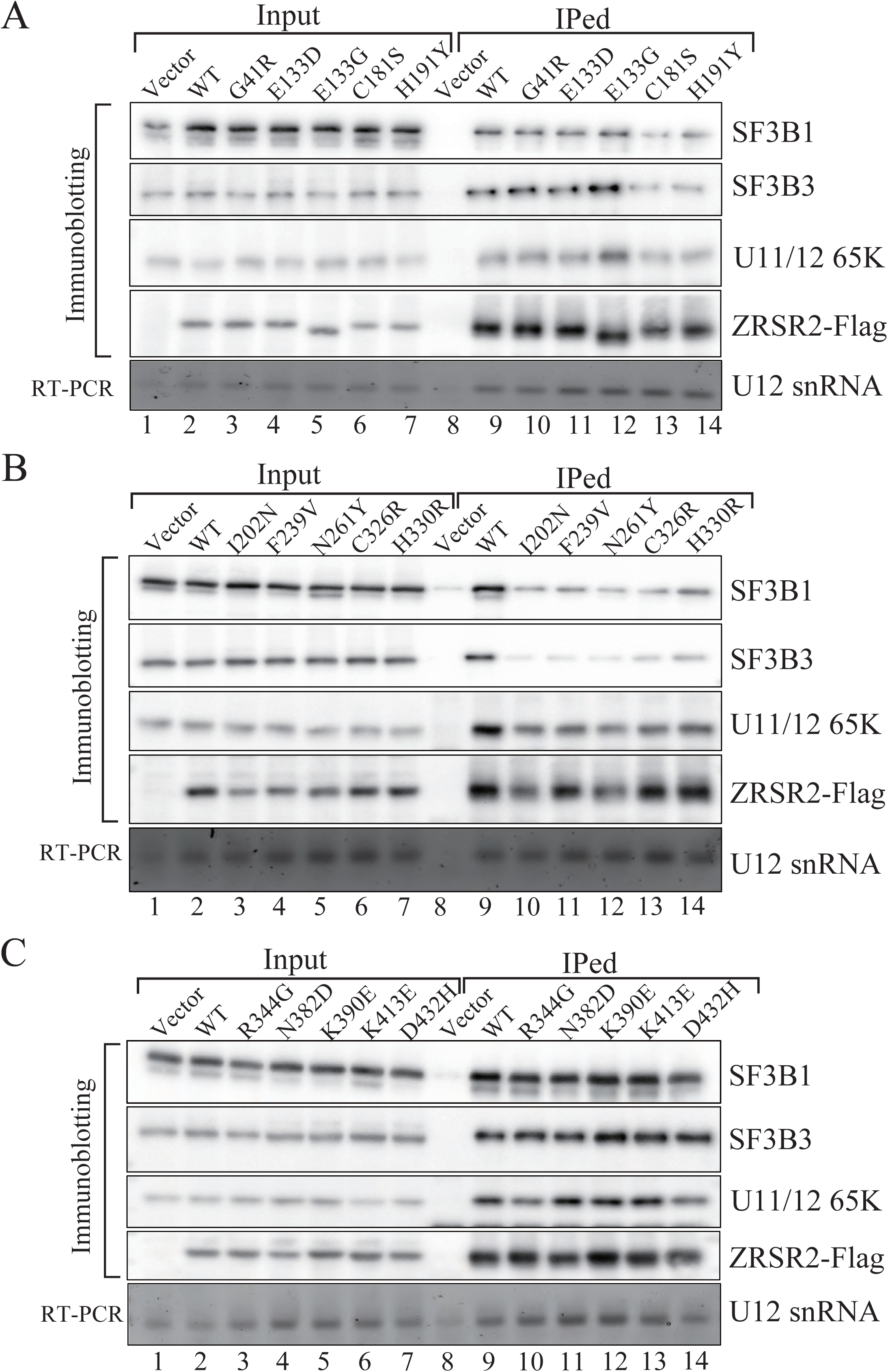
Co-immunoprecipitation of SF3B proteins and U11/12 65K protein with ZRSR2 mutant proteins found in MDS patients. Co-immunoprecipitation analysis utilizing wild-type or mutant ZRSR2 proteins that have missense mutations in the N-terminal or Zn1 domains (A), UHM or Zn2 domain (B), and BARD or RS domain (C). The co-precipitated proteins were analyzed by immunoblotting utilizing antibodies as indicated. Co-precipitation of U12 snRNA was also determined by RT-PCR.

### Mutations reducing interaction ability of ZRSR2 with SF3B impair the activity to support minor intron splicing

Lastly, to examine the effects of reduced interaction with SF3B on minor intron splicing mediated by ZRSR2, we carried out knockdown experiments by using shRNAs followed by adding back either wild-type or mutants of ZRSR2 protein. We established HEK293T cell lines stably expressing two different shRNAs targeting ZRSR2, and we found that both successfully reduced ZRSR2 mRNA levels (Figure 6A). We first utilized the NOP2/P120 minor intron splicing reporter to monitor the splicing supporting activity of ZRSR2, since wild-type ZRSR2 has been shown to facilitate the minor intron splicing of P120 minigene (Shen et al. 2010). The NOP2/P120 minor intron splicing reporter was co-transfected into ZRSR2-knockdown HEK293T#1 cells with either wild-type ZRSR2 or mutant ZRSR2 plasmids. As shown in Figure 6B, over-expression of wild-type ZRSR2 promoted splicing of P120 pre-mRNA (lane 2). In contrast, overexpression of C181S, F239V and C326R, which have a point mutation in Zn1, UHM and Zn2, respectively, exhibited less of a promoting effect on minor splicing than wild-type ZRSR2 (Figure 6B, lanes 4-6). These results demonstrate that the mutations in those three domains significantly reduce splicing supporting ability of ZRSR2. Overexpression of E133G (N-terminal domain), K390E (BARD) and D432H (RS domain) enhanced splicing of P120 pre-mRNA to a similar extent as wild-type (Figure 6B, lanes 7,8), indicating that these mutations do not affect ZRSR2 activity in minor intron splicing. We next analyzed the splicing supporting ability of ZRSR2 with endogenous minor introns. As splicing of minor introns in CLCN7 and USP21 genes has been reported to be affected in MDS patients who have mutations in ZRSR2 and those pre-mRNAs were cross-linked and immunoprecipitated with ZRSR2 (Madan et al. 2015; Inoue et al. 2021; Kwon et al. 2024), we employed these two genes and examined their splicing patterns under the overexpression of either ZRSR2 wild-type or mutant proteins. Results in Figure 6C showed that overexpression of wild-type ZRSR2 facilitated the minor intron splicing of CLCN7 but F239V and C326R mutants failed (Figure 6C, lanes 2-4). Similar results were obtained with USP21 pre-mRNA that the mutants tend to lose supporting activity for minor intron splicing (Figure 6D, lanes 2-4). Taken together, these results indicate that UHM and Zn2 mutations abolish the supporting ability of ZRSR2 in minor intron splicing by reducing its interaction with SF3B proteins.

**Figure 6.**
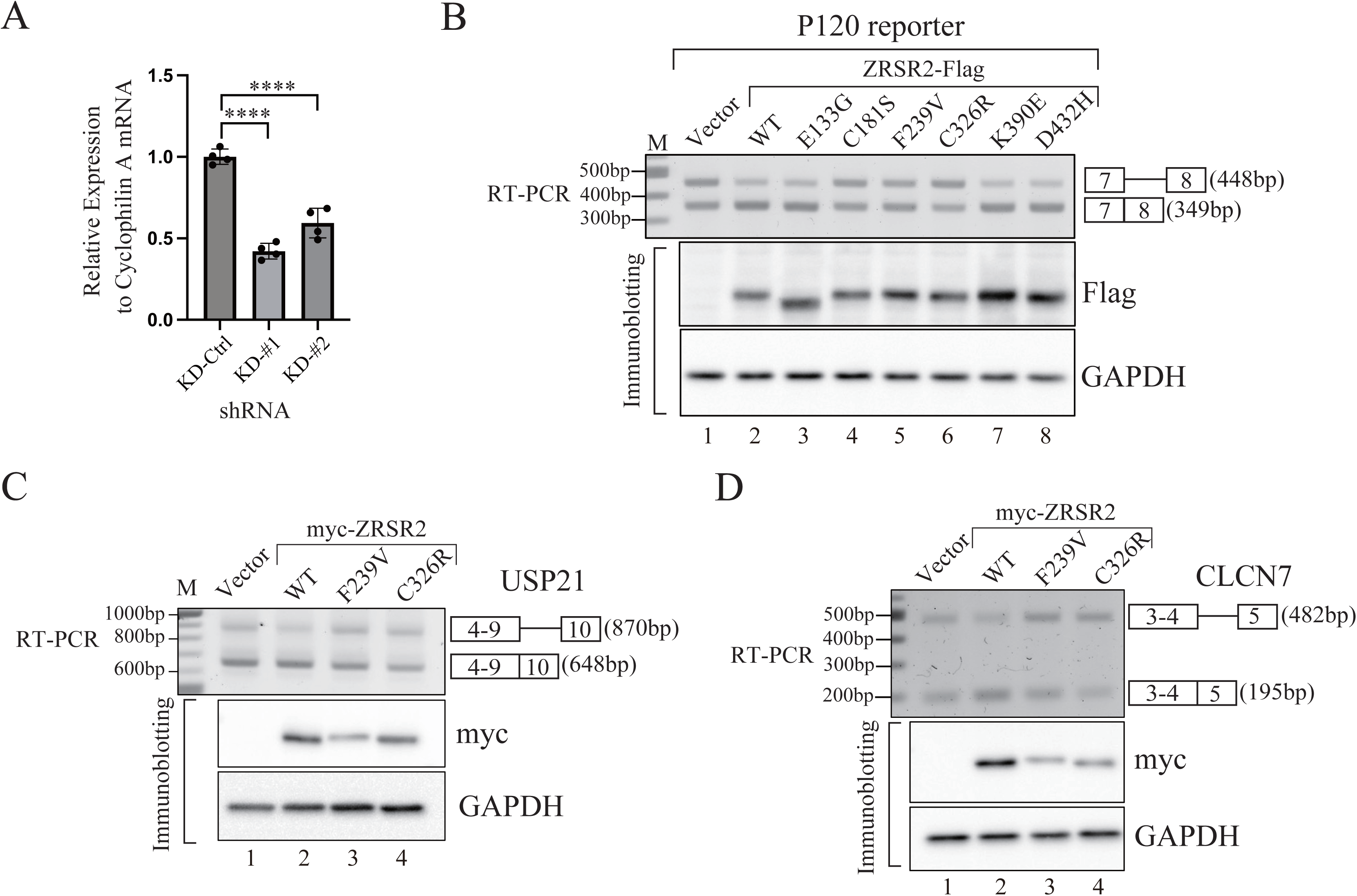
Mutations of ZRSR2 in Zn1, UHM and Zn2 found in MDS patients cannot support ZRSR2-dependent minor intron splicing. (A) Knockdown efficiency of ZRSR2 by shRNAs in HEK293T cells. The relative expression level of ZRSR2 mRNA is quantified by quantitative RT-PCR and normalized to Cyclophilin A mRNA. Ctrl: negative control shRNA targeting GFP. **** p<0.0001 (Dunnett’s multiple comparisons test) (B) Splicing assay of exogenous P120 reporter under overexpression of either ZRSR2 wild-type or mutant proteins. Plasmids expressing Flag-tagged ZRSR2 wild-type or mutants and P120 reporter were co-transfected into ZRSR2 knockdown HEK293T #1 cells and the splicing pattern of P120 reporter was determined by RT-PCR. Expression of ZRSR2 proteins was analyzed by immunoblotting with the anti-FLAG-tag antibody and GAPDH was detected as a loading control. (C, D) (C) Splicing assay of endogenous minor intron under overexpression of either ZRSR2 wild-type or mutant proteins. The myc-tagged ZRSR2 wild-type or mutant proteins were expressed in ZRSR2 knockdown HEK293T #1 cells and splicing pattern of minor intron in USP21 pre-mRNA (C) or CLCN7 pre-mRNA (D) was determined by RT-PCR. Expression of ZRSR2 proteins was analyzed by immunoblotting with the anti-myc tag and GAPDH was detected as a loading control.

## Discussion

In this paper we utilized ZRSR2 proteins carrying point mutations identified in MDS patients. Co-immunoprecipitation experiments identified SF3B proteins that were precipitated with wild-type ZRSR2 but not efficiently with UHM mutants (Figure 2). It is highly likely that this interaction determines subnuclear localization of ZRSR2, since SF3B binding deficient mutants exhibit different subnuclear localization from that of wild-type ZRSR2 (Figure 1B). The ZRSR2-SF3B interaction is mediated through UHM-ULM binding, like U2AF2-SF3B1 interaction (Thickman et al. 2006; Loerch and Kielkopf 2016; Galardi et al. 2022) (Figure 4). Mutations in Zn1 and Zn2 domains also reduced the interaction with SF3B (Figure 5). One possible mechanism is that those mutations affect the structure of Zn1/2 domains, and consequently overall structure of Zn1-UHM-Zn2 domains is altered. Structural analysis with SF3B1 ULM domain and ZRSR2 Zn1-UHM-Zn2 domains will validate this hypothesis.

Both wild-type ZRSR2 and SF3B-binding deficient mutants were able to co-precipitate U12 snRNA and U11/12 65K protein (Figure 5). Structural analysis of assembled minor intron spliceosome reveals that ZRSR2 is in the early spliceosome (Bai et al. 2024). One possible role of ZRSR2 during splicing is to recruit the SF3B complex to pre-mRNA, with which ZRSR2 and U11/12 snRNPs already associate. If this recruitment does not take place properly with some minor introns, these introns are not spliced out but are retained. We think this is the molecular mechanism of the assumed ‘loss of function’ caused by ZRSR2 mutations in MDS patients. Currently it is not known why ZRSR2 preferentially associates with SF3B complex, but not with SF3A complex. As we demonstrated, ZRSR2 more efficiently co-precipitates U11/U12 65K protein than U2A (Figure 3A). We assume that ZRSR2 binds to U11/U12 65K protein with higher affinity than to U2A protein, which in turn recruits U12 snRNP. Then it is possible that U12 snRNP-ZRSR2 association recruits only SF3B complex and avoids SF3A complex interaction with them, since U12-mediated minor intron splicing requires only SF3B complex (Turunen et al. 2013). Further biochemical experiments including reconstitution of this complex are required to test this hypothesis.

We have tested many ZRSR2 mutants that harbor point mutations in each domain. As described above, mutations in Zn1, UHM and Zn2 domains reduce interaction with SF3B complex (Figure 5A, B), which results in aberrant splicing of some minor introns (Figure 6B, C, D). However, other MDS-causing mutations in N-terminal domain, BARD and RS domain did not affect interaction of ZRSR2 with SF3B proteins (Figure 5A, C). In addition, none of the mutant proteins we tested showed reduction of the amount of co-precipitated U12 snRNP (Figure 5A, C). We also employed some mutants (E133G, K390E and D432H) for splicing assays with P120 splicing reporter (Figure 6B), but they showed splicing activity like that of wild-type ZRSR2 protein. These results indicate that such missense mutations may disrupt functions of ZRSR2 other than minor intron splicing. A recent study conducted by Fu et al. suggests other potential functions of ZRSR2 by showing that ZRSR1, a paralogous protein to ZRSR2 is involved in transcription regulation (Fu et al. 2026). Further comprehensive analysis into ZRSR2 functions is required to determine impacts of other MDS-causing mutations.

Although we found that UHM and Zn1/2 domains contribute to its binding to SF3B1, the functions of other domains of ZRSR2, such as N-terminal domain, BARD and RS domain, are not clarified yet. As for interaction with U12 snRNP, all the mutants we tested could co-precipitate U12 snRNA and U11/12 65K protein. Therefore, it is not known which domain mediates this interaction. In addition, it is not revealed which domain of ZRSR2 confers RNA binding activity. Specific RNA sequences that ZRSR2 binds are not determined, either. Functional domain structure analyses of ZRSR2 will shed light on the functions of each ZRSR2 domain.

## Materials and Methods

### Plasmid Construction

pcDNA3.1(+)-Flag plasmid containing Flag-tag coding sequence followed by stop codon (TAA) between XbaI and DraII sites in pcDNA3.1 (+) (Invitrogen) was a kind gift from Dr. Fumihiko Hakuno (The University of Tokyo). To construct C-terminal Flag-tagged ZRSR2 expressing vector (ZRSR2-Flag plasmid), the human ZRSR2 coding region was amplified by PCR using the following primers (ZRSR2_Full_F and ZRSR2_Full_R listed in Supplemental Table S1). Amplified products were cloned into pcDNA3.1(+)-Flag between BamHI and XhoI sites. Mutant ZRSR2-expressing vector and shRNA resistant ZRSR2 expressing plasmid were generated using Prime STAR Mutagenesis Basal Kit (TaKaRa Bio) with the ZRSR2-Flag plasmid as a template. Similarly, to construct N-terminal Flag -tagged or myc-tagged ZRSR2 expressing vector (Flag -ZRSR2 plasmid or myc-ZRSR2 plasmids), amplified ZRSR2 cDNA products were cloned into Flag-pcDNA3 or myc-pcDNA3 (Masaki et al. 2015). Mutant ZRSR2 expressing vector and shRNA resistant ZRSR2 expressing plasmid were generated by Prime STAR Mutagenesis Basal Kit (TaKaRa Bio) using Flag-ZRSR2 or myc-ZRSR2 plasmid as a template. To construct P120 minigene reporter, the human NOP2/P120 genomic DNA fragment containing exon7-intron7-exon8 was amplified by PCR and cloned into pcDNA3.1(+) vector using In-Fusion HD Cloning Kit (TaKaRa Bio).

For the GST-pull down assay, the cDNA fragments encoding ZRSR2 amino-acid residues 1-356 and 151-356 were amplified by PCR using cDNA. The products were cloned into the pET28b vector, in which an SD sequence, an initial ATG, a hexa-histidine tag, and a tobacco etch virus (TEV) protease cleavage site had been cloned immediately upstream of the protein coding region (pET28-His-TEV). cDNAs encoding SF3B1 amino acid residues 330-394, 330-349, and 348-394 were cloned into another modified pET28b vector, pET-GST-TEV, in which the GST-coding sequence had been inserted instead of the His-tag in pET-His-TEV.

### Cell Culture and Transfection

HeLa, HEK293T and HEK293TN cells were cultured in D-MEM (High Glucose) with L-Glutamine, Phenol Red and Sodium Pyruvate (Wako) supplemented with 10% fetal bovine serum (FBS; Gibco) and 1x Antibiotic-Antimycotic Mixed stock (Nacalai Tesque). Cells were grown in a 37℃ humidified atmosphere of 5% CO2. Transfection was carried out by using polyethylenimine (PEI) or TransIT-X2 Dynamic Delivery System (TaKaRa Bio) or Lipofectamine 2000 (Invitrogen) following the manufacturers’ recommendation. ZRSR2 knockdown HEK293T cell lines were generated by following the method as previously described (Madan et al. 2015). Lentiviral transduction was performed following the method as previously described (Okuda et al. 2022). Lentiviral shRNA vector (pLKO.1, Addgene) plasmids containing short hairpin RNA (shRNA) sequences targeting ZRSR2 sh1 (5’-CAACAGTTCCTAGACTTCTAT-3’), ZRSR2 sh2 (5’-AGCAGCCCTTTCTCTGTTTAA-3’) and GFP as a control (5’-GTCGAGCTGGACGGCGACGTA-3’) were transfected along with pMDLg/pRRE, pRSV-rev, and pMD2.G vectors (Dull et al. 1998) into HEK293TN cells using PEI. After 48 hours, the supernatant containing lentiviral particles was collected, centrifuged at 48.1 x g for 5 min and filtered. HEK293T cells were transduced with the prepared lentiviral particles in the presence of polybrene (8 μg/ml) for 24 h. Transduced cells were selected with 1.2 μg/ml puromycin to generate stable knockdown cell lines. The knockdown efficiency was verified by quantitative real-time PCR analysis using the same primer sets as used previously (Madan et al. 2015).

### Co-Immunoprecipitation (co-IP) assay

HEK293T cells transfected with ZRSR2-expressing plasmids were harvested in NET2 buffer (150 mM NaCl, 0.05% NP-40, 50 mM Tris-HCl (pH 7.5)) (Fukumura et al. 2023) or Buffer E (20 mM Tris-HCl (pH 8.0), 100 mM KCl, 0.2 mM EDTA, 0.2 mM PMSF) (Masaki et al. 2015). Cell lysates were sonicated using a VP-15S Ultras Homogenizer (TAITEC) and centrifuged for 10 min at 17800 × g at 4 °C. The supernatants were incubated with ANTI-Flag M2 Affinity Gel (Sigma) for 1 hour at 4 °C with gentle rotation. In the RNase treatment assay, 0.1 mg/ml RNase A (NIPPON GENE) was added to the IP mixture. After the 1 hour rotation, the beads were washed with NET2 or Buffer E for five times. For protein extraction, beads were suspended in the Sample Buffer Solution with Reducing Reagent (6x) for SDS-PAGE (Nacalai Tesque). Proteins were analyzed by SDS-PAGE followed by immunoblotting or silver staining. For RNA isolation, the beads were suspended in the Phenol:Chloroform:Isoamyl Alcohol 25:24:1 Mixed, pH 7.9 (PCI) solution (Nacalai Tesque), followed by ethanol precipitation.

### Mass spectrometric identification of proteins

Mass spectrometric analysis was carried out as previously described (Masaki et al. 2015). After separation by SDS-PAGE, the proteins were visualized by silver staining. The excised proteins were digested and subjected to matrix-assisted laser desorption/ionization time-of-flight mass spectrometry (MALDI-TOF/MS) by Thermo Fisher Scientific. Proteins were identified by comparing the molecular weights determined by MALDI-TOF/MS and the theoretical peptide masses of proteins registered in the NCBInr database using MASCOT software (Matrix Science, London, UK; http://www.matrixscience.com/search_form_select.html).

### GST Pull-down assay

His-tagged ZRSR2 and GST-fused SF3B1 expression plasmids were simultaneously transformed into the BL21(DE3) RIPL CodonPlus *E. coli* strain (Stratagene). The proteins were expressed in LB medium overnight at 15 °C after induction with 0.5 mM IPTG. His-tagged ZRSR2 and GST-fused SF3B1 were purified using Ni-NTA agarose (Qiagen) and glutathione Sepharose (GE Healthcare), respectively.

2 nmol GST-tagged SF3B1 was incubated with 4 nmol His-tagged ZRSR2 at room temperature overnight in reconstitution buffer containing 20 mM Tris/HCl (pH 8.0), 150 mM NaCl. 20 µl glutathione Sepharose resin was used for the pull-down assay. After washing by repeated centrifugation and resuspension in the reconstitution buffer, the complex was eluted by 50 µl elution buffer (20 mM reduced glutathione in the reconstitution buffer). Proteins were separated by SDS–PAGE (15 %) and stained with Coomassie brilliant blue.

### Minigene or endogenous minor intron splicing assay

For the reporter splicing assay, we co-transfected a P120 minigene reporter and shRNA-resistant ZRSR2-expressing plasmids into ZRSR2-knockdown HEK293T cells (KD#1). In the endogenous minor intron splicing assay, only the shRNA-resistant ZRSR2 expression vector was transfected into ZRSR2-knockdown HEK293T cells (KD#1). After 48 hours of transfection, total RNA was isolated using Trizol Reagent (Molecular Research Center) according to the manufacturer’s instructions. Proteins were extracted by harvesting cell pellets in Lysis Buffer (25 mM Tris-HCl (pH 7.5), 150 mM NaCl, 0.1 % SDS, 1 % NP-40, 0.5 % Sodium Deoxycholate). After freezing and thawing cell lysates, followed by a centrifugation for 10 min at 17800 x g (4 °C), the supernatant fractions were mixed with the Sample Buffer Solution with Reducing Reagent(6X) for SDS-PAGE (Nacalai Tesque).

### Reverse transcription–PCR (RT–PCR) analysis

Isolated RNA was reverse transcribed using ReverTra Ace^®^ qPCR RT Master Mix with gDNA Remover (TOYOBO). Transcribed cDNA was used as a template for conventional PCR analysis (SapphireAmp^®^ Fast PCR Master Mix, TaKaRa Bio) or quantitative real-time PCR (qPCR) analysis (KAPA SYBR FAST qPCRMaster Mix (2X) Universal, Nippon Genetics). PCR primers used in this study are listed in Supplemental Table S1.

### Immunofluorescence assay

HeLa cells transfected with C-terminal Flag-tagged ZRSR2 plasmids were cultured on coverslips, fixed with 4% paraformaldehyde, permeabilized with 0.5% Triton-X-100/PBS (-), and blocked with 5% BSA/PBS (-). Primary antibody against Flag-tag (Anti-DDDDK-tag mAb, M185-3L, MBL) and the FITC-conjugated AffniPure Donkey α-mouse IgG (H+L) (Jackson Immuno Research Laboratories, Inc.) as secondary antibody were used. Nuclei were stained using Hoechst 33342 solution (5μg/ml, Sigma). Coverslips were mounted with Vecta-shield mounting medium (VECTOR Laboratories) on glass slides for fluorescence microscopy using BZ-8000 (KEYENCE).

### Immunoblotting analysis

The samples were separated on SDS-PAGE and transferred to Immobilon PVDF membranes (Millipore, Burlington) by electroblotting. After blocking, the membranes were incubated with primary antibodies for the following proteins: SAP120/SF3A1 (204 011, Synaptic Systems), SAP155/SF3B1 (D221-3, MBL), SAP130/SF3B3 (ab3608, Abcam), SAP49/SF3B4 (ab183483, Abcam), U2 snRNP A (sc-393804, Santa-Cruz) U11/12 snRNP 65K/RNPC3 (sc-514951, Santa-Cruz), Flag-tag (M185-3L, MBL or F7425-.2MG, SIGMA), myc-tag (9E10, Santa-Cruz), GAPDH (#2118, Cell Signaling) The bound antibodies were detected with peroxidase-conjugated secondary antibodies (Jackson ImmunoResearch Laboratories) and signals were developed with Chemi-Lumi One Super detection reagents (Nacalai Tesque). The signals were detected by GeneGnome System High Resolution 100 (SYNGENE) or WSE-6170-CP Lumino G (ATTO).

## Acknowledgements

We thank Dr. Fumihiko Hakuno for a kind gift of pCDNA3.1-Flag plasmid. We are grateful to the members of Laboratory of Cellular Biochemistry for critical discussion.

## Author Contribution

Tomoki Chiba: Formal analysis, Investigation, Methodology, Visualization, Writing -original draft; Eri Matsumoto: Investigation, Methodology, review & editing; Eiji Obayashi: Resources, Investigation, Writing -review & editing; So Masaki: Formal analysis, Investigation, Writing ‒ review & editing; Satoshi Tanaka: Resources, Writing -review & editing; Naoyuki Kataoka: Conceptualization, Funding acquisition, Project administration, Supervision, Resources, Writing -original draft.

