## Supplemental Table 1 for "MDS-causing mutations of ZRSR2 impair minor intron splicing by reducing its interaction with SF3B complex"

Supplemental Table S1. Primers sequences used in this sutdy

| Name | Sequence (5' to 3') | Application |
|---|---|---|
| P120_ex7_F | GCTGGGGAGATGGAGCAG | RT-PCR |
| pcDNA3.1_R | CTAGAAGGCACAGTCGAGGCT | RT-PCR |
| CLCN7_ex3.4_F | ATCAATCACACGGCCTTCCG | RT-PCR |
| CLCN7_ex5_R | AGCAACAGGGAGAAGGACAG | RT-PCR |
| USP21_ex4_F | CCTACTCGATTCCGAGCTGTC | RT-PCR |
| USP21_ex10_R | GTCACTGGCAAAGTCCCCTA | RT-PCR |
| U12snRNA_F | AACTTATGAGTAAGGAAAATAACGATTCG | RT-PCR |
| U12snRNA_R | CGACCTTTACCCGCTCAAAA | RT-PCR |
| hZRSR2_F | AAGATGACGTTTCCCGAGAAACC | qRT-PCR |
| hZRSR2_R | ATAAAAGTGTCCTCCTCTTCCTCCTT | qRT-PCR |
| hCyclophillineA_F | GTCAACCCCACCGTGTTCTTC | qRT-PCR |
| hCyclophillineA_R | TTTCTGCTGTCTTTGGGACCTTG | qRT-PCR |
| ZRSR2_Full_F | AAAGGATCCATGGCTGCGCCCGAGAAGATGACG | Cloning |
| ZRSR2_Full_R | AAACTCGAGTTTGGATTTGGGACTCTGAACAGT | Cloning |
| P120reporter_F | GTTTAAACTTAAGCTTGATCCAGTGGAGTGAAGAG | Cloning |
| P120reporter_R | GCTCGGTACCAAGCTTCTCAGACAGAGGGAAGAG | Cloning |
